# Identification and expression analysis of Shaker K^+^ channel genes in millet *(Setaria italica)* under abiotic stresses

**DOI:** 10.1101/2021.09.30.462669

**Authors:** Ben Zhang, Hui Wang, Yue Guo, Xiaoxia Wang, Mengtao Lv, Pu Yang, Lizhen Zhang

## Abstract

Potassium (K+) is one of the essential nutrients for plant, which is involved in plant growth and development and abiotic stress tolerance. The absorption and transport of K^+^ depends on Shaker K+ channels. Foxtail millet is a Poaceae crop with strong drought stress-tolerant. In this study, we identified ten Shaker K^+^ channel genes in foxtail millet. Phylogenetic analysis, prediction of conserved motif, and gene structure analysis classified these genes into five groups. The transcription level of these genes under different abiotic stress treatments (cold, heat, NaCl, PEG) and ABA treatment were analyzed by quantitative real-time PCR. Each gene displayed its own regulation pattern under different treatments, suggests these channels play important role in plant adaptation to different environment conditions.

## Background

Drought, salt, and temperature stresses as the main abiotic stresses severely affect plant growth and development. Potassium (K^+^) is one of the essential nutrients for plant, which is involved in most physiological processes of plant growth and development (Ward et al., 2009). K^+^ is involved in plant cell turgor maintenance, stomata movement, photosynthesis, carbon assimilation, and osmotic balance, which suggest its role in plant abiotic stress tolerance (Wang et al., 2013). Although many reports have suggested the protective role of K^+^ under abiotic stress in different species (Ma et al., 2020; Gong et al., 2020), the molecular mechanism of K^+^ in plant abiotic stress tolerance is still not well understand.

In plant, there are two main mechanisms for the absorption of K^+^, namely the high-affinity K^+^ absorption mechanism mediated by transporters and the low-affinity K^+^ absorption mechanism mediated by potassium channels (Raddatz et al., 2020). The high affinity K^+^ transporters in millet have been analyzed (Zhang et al., 2018). However, there is little report focused on K^+^ channels in millet. The K^+^ channels are divided into three families according to their α-subunits configuration differences, including Shaker, TPK and Kir-like K^+^ channel families (Lebaudy et al., 2007). Among them, the Shaker K^+^ channel family is the most extensively studied and has been identified in many plants, including Arabidopsis (Dreyer and Uozumi, 2011), rice (Amrutha et al., 2007; Hwang et al., 2013), maize (Bauer et al., 2000), poplar (Zhang et al., 2010), and sweetpotato (Jin et al., 2021). Shaker K^+^ channel have been shown to be involved in K^+^ uptake from soil and stomata movement. Their subunits share similar common structural features. They all contain six transmembrane α-helices, called S1-S6, and every four subunits form a functional channel as tetrameric assemblies, which mediates the transport of potassium ions across the membrane (Dreyer and Blatt, 2009). In Arabidopsis, the nine members of Shaker K^+^ channel genes can be further divided into five groups (Pilot et al., 2003b). Group I (KATs) and Group II (AKT1, AKT5, AKT6) are inward rectification channel, whose open are regulated by the negative membrane voltage. By contrast, Group V (SKOR and GORK) is the outward rectification channel, whose open requires the membrane depolarize to positive voltages. The Group III (AKT2) belongs to the weak inward rectification channel and Group IV contains a Silent subunit KC1, and Group V (SKOR and GORK) is outward rectification channels (Lebaudy et al., 2007). A functional channel are homo- or hetero-tetrameric assemblies of proteins from these groups (Jeanguenin et al., 2008; Duby et al., 2008).

The role of Shaker K^+^ channels in plant abiotic stress response has been investigated. The cold and salt stress induced expression of GORK, leading to the disruption of cytosolic Na^+^/K^+^ ratio and suppression of plant K^+^dependent metabolism (Chen et al., 2007; Becker et al., 2003). For drought stress, most of Shaker K^+^ channels, including KAT1, KAT2, AKT1, AKT2, KC1, and GORK, are known to be involved in opening and closing of stomata (Hosy et al., 2003; Lebaudy et al., 2008). The K^+^ efflux from guard cell through GORK and the channel subcellular location are regulated by drought stress related phytohormone abscisic acid (ABA) through an OPEN STOMATA 1 (OST1) kinase involved signaling system (Chen et al., 2021). Ooi et al. have found there was direct GORK–ABA interaction which increased the GORK K^+^current (Ooi et al., 2017). Overexpressing inward rectification channel OsAKT1 in rice improved the tissue K^+^ content and plant drought tolerant (Ahmad et al., 2016). Similar function also reported for overexpressing HvAKT1 in barley (Feng et al., 2020). The inward rectification channel KAT1 is also regulated by ABA (Sutter et al., 2007). Recent reports suggest such ABA related K^+^ channel regulation depends on the interaction between KAT1 and membrane traffic related SNARE family protein SYP121 (Sutter et al., 2006; Honsbein et al., 2009). SNARE proteins help to drive membrane fusion during vesicle traffic and involved in abiotic stress response (Bassham and Blatt, 2008; Salinas-cornejo et al., 2019; Zhang et al., 2020). Further research found that VAMP721, the partner of SYP121 during membrane fusion, also binds with K^+^ channels but regulates the channel activity in opposite way (Zhang et al., 2015), suggesting there is a SNARE proteins related Shaker K^+^ channel regulation system. The expression of SNARE proteins are induced by abiotic stress (Wang et al., 2021; Kwon et al., 2020). Thus, the SNARE related regulation of K^+^ channels might be the key to explain the mechanism of K^+^-dependent plant abiotic tolerance.

*Foxtail millet* is a Poaceae crop with strong environmental adaptability. The high abiotic stress tolerance of millet leads to its wildly cultivated in Asia and Africa (Peng and Zhang, 2020; He et al., 2015). In 2012, the genome of millet was sequenced, making it possible to identify genes from millet and study their functions (Bennetzen et al., 2012; Zhang et al., 2012). Furthermore, its relatively small genome makes millet as a promising C4 model plant (He et al., 2015). Previous research from our group has analyzed the role of SNARE proteins under drought stress in millet (Wang et al., 2021). However, there is limited report focused on Shaker K^+^ channels and their role in abiotic stress tolerant in millet. Therefore, based on the sequence information of Shaker K^+^ channel genes from Arabidopsis and rice, here, we identified the Shaker K^+^ channel genes in the millet database, and finally obtained 10 *Shaker K^+^ channel* gene members. Furthermore, these genes were further analyzed from the aspects of phylogeny, conserved motifs and domain, promoter, and tissue expression pattern. At the same time, quantitative real-time PCR (qPCR) was used to analyze the transcription levels of *Shaker K^+^ channel* family genes in foxtail millet under different abiotic stresses. This is conducive to exploring the mechanism of K^+^ absorption in millet in response to abiotic stress, and provides directions for future millet genetic transformation and physiological research. In addition, we also analyzed whether there was similar interaction between Shaker K^+^ channel protein and SNARE proteins in C4 plant millet as previous reports in Arabidopsis, which is beneficial to realize the combination of theory research in model plant and application research in crop.

## Methods

### Identification of *Shaker K^+^ channel* genes in millet

The Shaker K^+^ channel sequences of Arabidopsis and rice were extracted from the Arabidopsis Information Resource (http://www.arabidopsis.org/) and the Rice Genome Annotation Project (http://rice.plantbiology.msu.edu/index.shtml). These sequences were used to blast in the Phytozome database (Phytozome 13, Setaria italica v2.2; https://phytozome-next.jgi.doe.gov/) (Goodstein et al., 2012) to obain the candidate shaker K^+^channel coding sequences. Then the Arabidopsis channel sequence was submitted to the Pfam database (http://pfam.xfam.org/) to obtain the hidden Markov model (HMM) profile of the Shaker K^+^ channel domain, which were queried against the candidate sequences of shaker K^+^channels from millet. Finally, a sequence with a complete channel conserved domain was selected for subsequent analysis. Based on the protein sequence, the theoretical isoelectric point (PI), molecular weight (MW), and amino acid composition of the Shaker K^+^ channels from millet were analyzed by ExPASY (http://web.expasy.org/protparam)(Gasteiger et al., 2003). Their sub-cellular localization was predicted by WoLF PSORT (https://wolfpsort.hgc.jp/) (Horton et al., 2007) and CELLO v2.5 Server (http://cello.life.nctu.edu.tw/) (Yu et al., 2004).

### Phylogenetic analysis and chromosome location of Shaker K^+^ channel family

The amino acid sequences of Shaker K^+^ channels from millet, arabidopsis (Cao et al., 1995), and rice (Hwang et al., 2013) were homologously aligned by the ClustalX software (Thompson et al., 2003) with default parameters. Based on the results of this comparison, MEGA7.0 (Kumar et al., 2016) software was used to construct the phylogenetic tree of the Shaker K^+^ channels with the following parameters: poission model and pairwise deletion, Bootstrap 1000 repetitions. The gene annotations of identified Shaker K^+^ channel genes were extracted from the Phytozome database (Phytozome 13, Setaria italica v2.2; https://phytozome-next.jgi.doe.gov/)(Goodstein et al., 2012), from which the position of the genes on chromosome were obtained, and the physical map is drawn using MapInspect software (https://mapinspect.software.informer.com/) (Wu et al., 2019).

### Motif Composition and Gene Structure Analysis of Millet *Shaker K^+^ Channel* genes Family

The protein conserved motifs of Shaker K^+^ channels from millet were predicted by online Multiple Em for Motif Elicitation (MEME) programme (http://meme.nbcr.net/meme/) (Bailey et al., 2009), with the following parameters: the number of repetition = any, the maximum number of motifs = 10. The CDs sequences and genome sequences of these genes were downloaded from the Phytozome database and submitted to Gene Structure Display Server online program (GSDS: http://gsds.cbi.pku.edu.cn) (Hu et al., 2015) for gene structure analysis.

### Cis-acting element analyses

To obtain the cis-elements in the promoter regions of Shaker K^+^ channels from millet, the 2000 bp upstream sequence of each genes were extracted from the Phytozome database and submitted to the PlantCARE software (http://bioinformatics.psb.ugent.be/webtools/plantcare/html/; Lescot et al., 2002).

### Tissue expression patterns in millet Shaker K^+^ channel family

In order to study the potential expression pattern of Shaker K^+^ channels from millet at different tissues and developmental stages, the fragments per kilobase of the exon model per million mapped (FPKM) values of these genes were obtained from the NCBI GEO RNA-seq DataSets (GSE89855). Including roots (at inflorescence stage), shoot (at inflorescence and vegetative stages), leaf (four locations: tip, midsection, lower section, and base at the vegetative stage), meristem (at inflorescence and vegetative stages), and panicle (at pre-anthesis and postanthesis stages). These data were submitted to TBtool (Chen et al., 2020) for expression profile mapping.

### Plant growth conditions and abiotic stress treatment

The foxtail millet variety “Jingu 21” was used for the treatment of different abiotic stresses in present study (seeds obtained from the stock of Prof. Lizhan Zhang’s lab, School of Life science, Shanxi University, China. plant materials were provided free of charge and used for research only). The plant seeds were grown in a tray containing vermiculite and nutrient soil at a ratio of 1:1 and cultivated under greenhouse conditions (16 h light/8 h dark at 23-26 °C, 50000 Lux light, and 30-50% relative humidity). When the seeds germinate and grow to the two-leaf stage (ten days), select seedlings with consistent growth were transferred to plastic pots with three plants per pot. For temperature stress, after another four days, millet seedlings were separately subjected to heat (40°C day/32°C night) and cold (4°C) stress in a constant temperature incubator, for 0, 12 and 24 h. For other abiotic stresses, ten days old millet seedlings were transferred to 1/2 MS liquid medium and cultured for another four days, then subjected to salt (150 mmol/l, 200 mmol/l), PEG6000 (10%,15%), and ABA (10 mmol/l) stress respectively, for 0, 12 and 24 h. There were three replicates for all stress treatments. The leave samples were harvested, immediately frozen in liquid nitrogen and stored at −80°C for further RNA extraction.

### RNA extraction and qRT-PCR

Total RNA is extracted from all experimental treatments using TransZolTM UP Plus RNA Kit (TransGen Biotech, Beijing, China) according to the manufacturer’s instructions. The first-strand cDNA templates were synthesized using EasyScript® One-Step gDNA Removal and cDNA Synthesis SuperMix Kit (TransGen Biotech, Beijing, China) in a total volume of 30 μl according to the manufacturer’s instructions. And the total volume of the qRT-PCR reaction system was 10 μl, which included 5 μl of TransStart® Tip Green qPCR SuperMix (TransGen Biotech, Beijing, China), 1 μl of diluted cDNA template, 0.8 μl of upstream and downstream primers (10μmol/l), and 3.2 μl of RNase free ddH_2_O. The qPCR thermal cycler program included 94 °C for 30 s; followed by 40 cycles at 94 °C for 15 s and 60 °C 30 s. ALL primers were synthesized by Shanghai Sangon Biotech (As shown in supplement Table 1), wherein *SiACT2 (Seita.8G043100)* was used as the internal reference. Each experiment includes three technical replicates and three biological replicates.

### Prediction of Potential N-Glycosylation and Phosphorylation Sites of Shaker K+ Channel Proteins

The sequences of Shaker K^+^ channel proteins from millet were submitted to NetOGlyc 4.0 (http://www.cbs.dtu.dk/services/NetOGlyc/) for N-glycosylation sites prediction, and NetPhos 3.1 (http://www.cbs.dtu.dk/services/NetPhos/) was used to predict protein phosphorylation sites (including Serine, Threonine, Tyrosine). The positive result was confirmed when the score >0.5.

### ABA and G protein binding site prediction in the millet

The presence of abscisic acid (ABA) interaction domain (D543 to G575) (Ooi et al., 2017) and guanine nucleotide–binding proteins (G-protein) binding motif (E165 to Y169)(Adem et al., 2020; Deupi et al., 2012) on the protein sequence of Arabidopsis AtGORK (At5g37500) have been reported. Due to the high similarity of GORK and SKOR sequences, in order to predict whether it will be similar binding sites exist in other plants, especially millet. Therefore, the GORK and SKOR amino-acid sequences of Arabidopsis (GORK, At5g37500; SKOR, At3G02850), tobacco (GORK, XP_016457756; SKOR, XP_016460239), millet (GORK, Seita.7G111600; SKOR, Seita.4G110300), rice (GORK, Os04g36740; SKOR, Os06g14030), sorghum (GORK, Sobic.006G093400; SKOR, Sobic.010G102800) and wheat (GORK, Traes_2BL_C71ACBCED; SKOR, Traes_7DS_965990A2F) were extracted from the Phytozome and NCBI database, and multiple sequence alignments were performed using ClustalX software.

## Results

### Identification of *Shaker K^+^ Channel* genes in foxtail millet

To identify the Shaker K^+^ channels in millet, the coding sequences of Shaker K^+^ channels from Arabidopsis and rice were used to blast in the Phytozome database, and 12 candidate genes were obtained. Then the Arabidopsis Shaker K^+^ channel protein sequences were submitted to the Pfam database (http://pfam.xfam.org/) to obtain the Shaker K^+^ channel domain (PF00520.31, PF00027.29, and PF11834.8). The protein sequences of these candidate genes were submitted to the SMART programme (http://smart.embl-heidelberg.de/) and finally identified that there were 10 Shaker K^+^ channel genes in millet (Table1). In addition, the encoded protein of the millet Shaker K^+^ channel genes were 505 (Seita.5G323200)-903 (Seita.2G049100) amino acids in length and molecular weight was distributed from 58.16-98.76 kDa. The theoretical isoelectric point calculation shows that the isoelectric point (pI) of Shaker K^+^ channel proteins range from 6.23 (Seita.4G110300) to 9.08 (Seita.1G210600). Predictions of subcellular localization indicated that all Shaker K^+^ channel proteins were localized on the plasma membrane (PM) with 4-6 trans-membrane segments (Table1).

**Table 1.** Detailed information of predicted Shaker K^+^ channel genes in Foxtail millet.

| Gene ID | Genome position |  | Chr | Length<br>(aa) | pI | Mw<br>(KDa) | Predicted<br>gene<br>name | Subcellular<br>localization | Number<br>of TM<br>segments | Orthologous<br>gene ID |
| --- | --- | --- | --- | --- | --- | --- | --- | --- | --- | --- |
|  | Start | End |  |  |  |  |  |  |  |  |
| Seita.1G020100 | 1728964 | 1735893 | chr1 | 761 | 7.6 | 87.45 | SiKAT3 | PM | 5 | AT2G25600,<br>AT4G18290,<br>AT4G32500,<br>AT5G46240,<br>Os02g14840 |
| Seita.5G244400 | 3068192 | 30688913 | chr5 | 885 | 6.66 | 99.66 | SiAKT1 | PM | 5 | AT2G26650,<br>Os01g45990 |
| Seita.4G110300 | 103449 | 10351307 | chr4 | 854 | 6.23 | 96.77 | SiSKOR | PM | 5 | AT3G02850,<br>Os06g14030,<br>AT5G37500 |
| Seita.3G233000 | 19292761 | 19297970 | chr3 | 855 | 8.58 | 94.53 | SiAKT2/3 | PM | 5 | AT4G22200,<br>Os05g35410 |
| Seita.5G298000 | 35360171 | 35362824 | chr5 | 617 | 8.92 | 69.5 | SiKC1a | PM | 6 | AT4G32650,<br>Os06g14310,<br>Os04g02720,<br>Os01g52070 |
| Seita.7G111600 | 21144981 | 21149199 | chr7 | 724 | 8.47 | 81.79 | SiGORK | PM | 5 | Os04g36740 |
| Seita.5G323200 | 37290994 | 37293812 | chr5 | 505 | 8.95 | 58.16 | SiKAT1 | PM | 5 | Os01g55200 |
| Seita.5G148000 | 1317289 | 13177173 | chr5 | 695 | 8.82 | 79.48 | SiKAT2 | PM | 4 | Os01g11250 |
| Seita.2G049100 | 3916917 | 3920473 | chr2 | 903 | 8.32 | 98.76 | SiAKT2 | PM | 5 | Os07g07910 |
| Seita.1G210600 | 28923321 | 28925051 | chr1 | 576 | 9.08 | 63.15 | SiKC1b | PM | 4 | Os06g14310 |

### Phylogeny analysis and Chromosome location Shaker K^+^ Channel genes in foxtail millet

To further study the evolutionary relationship of Shaker K^+^ channels between different species, the protein sequences from millet, Arabidopsis, and rice were used to construct a phylogenetic tree. As shown in Figure 1, the Shaker K^+^ channel proteins from millet can be classified into five groups based on their phylogenetic relationships and functional divergences as reported before (Pilot et al., 2003b; Hwang et al., 2013). The Group I and II were inward rectifying channels. There were SiAKT1, SiAKT2 in Group I and SiKAT1, SiKAT2, SiKAT3 in Group II. There was only SiAKT2/3 belong to Group III as a weakly inward rectifying channel. Group IV, the regulatory channel or silent channel, included SiKC1a and SiKC1b. Group V was outward rectifying channel included SiGORK and SiSKOR. Group consisted of Shaker K^+^ channels from different species suggested that the protein family is evolutionarily conserved. Compared to Arabidopsis, proteins from millet were more similar to their orthologous genes in rice. Interestingly, the channels in Group IV formed species-specific sub-groups. There were 2 in Group IV from millet, while the number in Arabidopsis and rice were 1 and 3.

**Figure 1.**
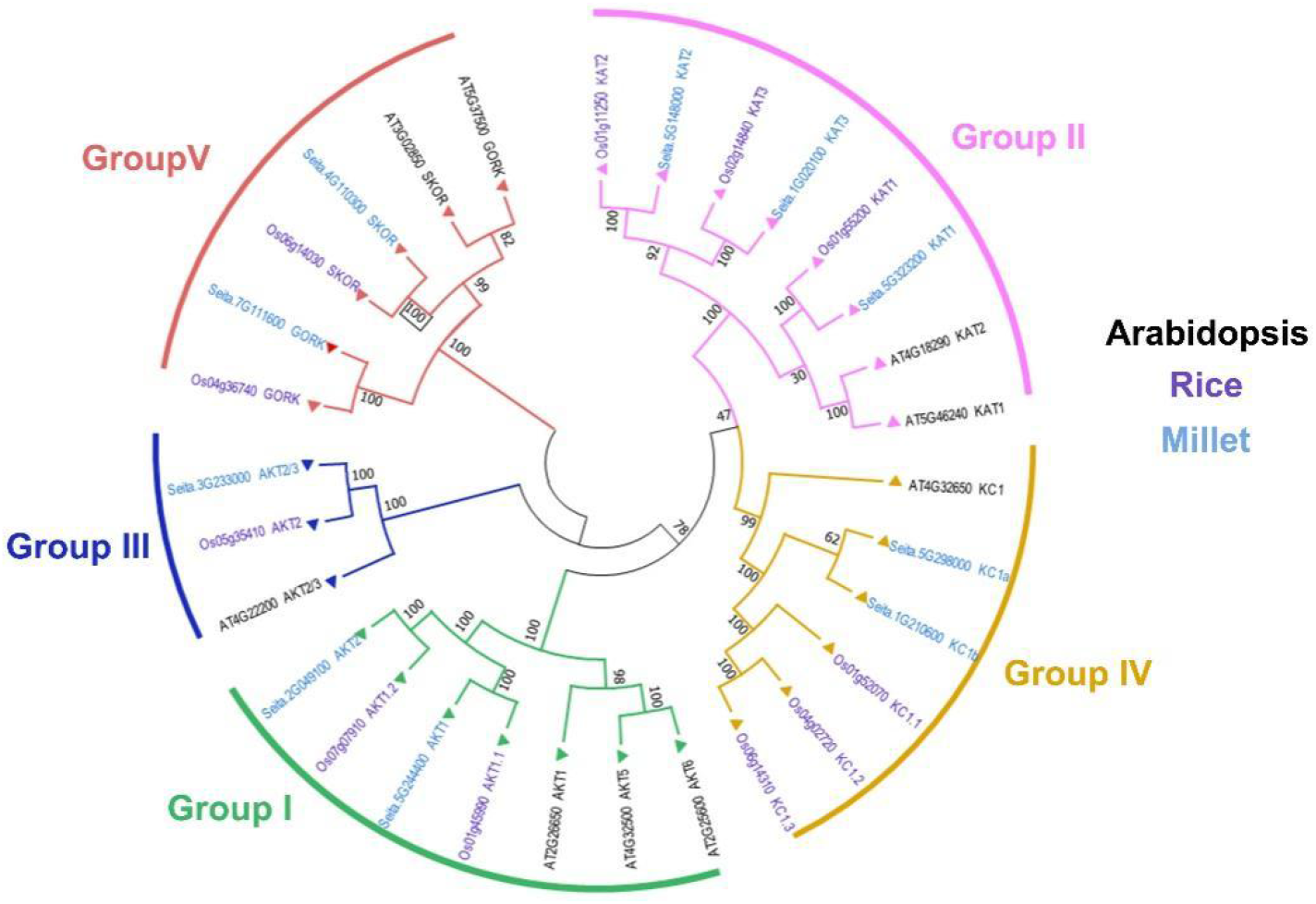
Phylogenetic analysis and classification of Shaker K^+^ channel proteins from millet (light blue), Arabidopsis (black), and rice (purple). Shaker K^+^ channel proteins are grouped into five distinct groups denoted by different colored lines.

### Chromosome location, gene structure, and motif analyses of Shaker K^+^ channel gene family

These Shaker K^+^ channel genes were distributed on 6 out of 9 chromosomes of millet (Figure 2). Chromosome 1 contained 2 Shaker K^+^ channel genes *(SiKAT3* and *SiKC1b)*, there was only one gene on Chromosome 2, 3, and 7. Chromosome 5 contained the greatest number of Shaker K^+^ channe*l* genes (4 genes).

**Figure 2.**
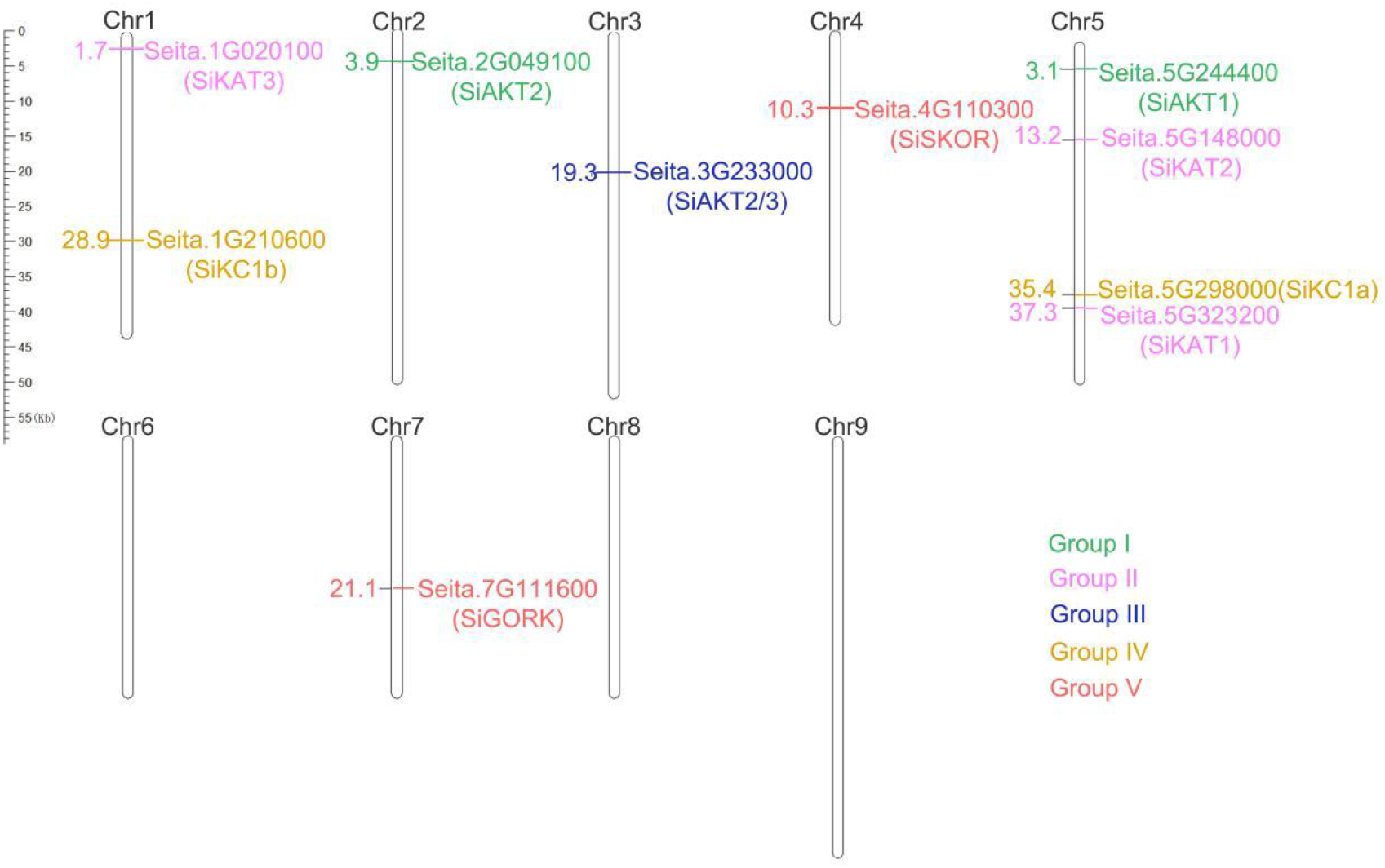
Chromosomal localization of Shaker K^+^ channel genes from millet. The location information of 10 *Shaker K^+^ channel* genes on the chromosome was obtained from the Phytozome database (Setaria italica v2.2). Different colours on the nine chromosomes (chr1-chr9) indicate different channel groups

Next, the gene structural diversity and protein motif distribution of the *Shaker K^+^ channel* genes in millet were explored. As shown in the right side of Figure 3, same group of genes contained similar numbers of introns and exons. For example, Group I, II and V contained 9-11 introns, and group III and IV contained 6-8 introns except for *SiKC1b* without introns. The differences in gene structure indicate the evolutionary and functional differences in the Shaker K^+^ channel family and provide additional evidence to support phylogenetic grouping.

**Figure 3.**
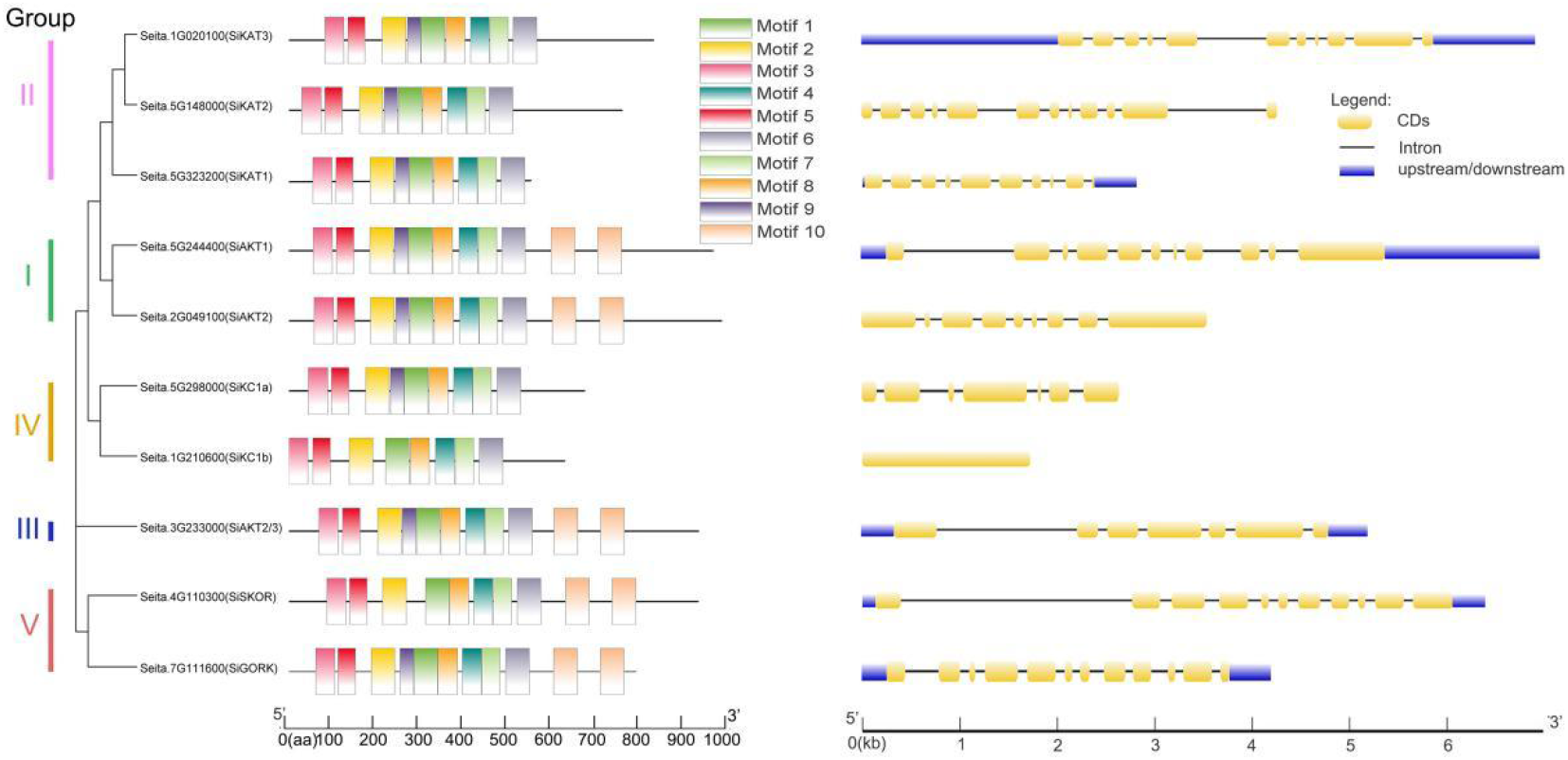
Protein motifs and gene structures of Shaker K^+^ channels from millet. Left: In the Shaker K^+^ channel proteins, 10 motifs were identified by the MEME tool, represented by different colours (1–10) and depicted by TBtools. Right: The exon–Intron structure of these *Shaker K^+^ channel* genes was predicted by GSDS 2.0. The yellow boxes represent the gene coding region (CDS), the black lines represent introns, and the blue boxes represented up/downstream untranslated Regions (UTR).

As shown in Figure 3 left, 10 conserved motifs were detected in Shaker K^+^ channel proteins by MEME. All channel proteins contained conserved motifs 1-8. Proteins in Group I, III, and V contained two motif 10. Motif 9 were missing from *SiSKOR* and *SiKC1b*. This analysis also supports our classification based on phylogenetic tree.

### Cis-acting regulatory elements analysis of *Shaker K^+^ channel* gene family

To further understand the regulatory mechanism the *Shaker K+ channel* genes, the 2000 bp promoter region upstream of them were used to analyze their cis-elements. As shown in Supplement Table 2, a total of 84 elements were identified. Based on the functional differences, important cis-acting elements were divided into three major classes: plant growth and development related, phyto-hormone responsiveness related, and abiotic/biotic stresses related cis-acting elements. As shown in Figure 4, circadian element (Anderson et al., 1994) was only found in *SiKC1a (Seita.5G298000)*, indicating that this gene may be related to photoperiod regulation. The RY elements are seed-specific promoters, which mediates initial transcriptional activation during embryo mid maturation (Lelievre et al., 1992;Guerriero et al., 2009). This cis-acting elements was only predicted in *SiKAT3 (Seita.1G020100)*. In addition, promoter elements related to plant growth, such as A-box, CAT-box and O2-site, were also found in 3, 6, and 6 *Shaker K^+^ channel* genes, respectively. ABA response elements (ABREs) (Hobo et al., 1999) was identified in 6 Shaker K+ channel genes. The *SiKAT1 (Seita.5G323200)* had 9 ABRE elements, *while SiAKT2/3 (Seita.3G233000)* had 8. The MeJA-responsive element CGTCA-motif and TGACG-motif (Wang et al., 2019) were found in 9 genes. The *SiAKT1 (Seita.5G244400)* had the largest number of these two elements (6 for each). Other phytohormone-related cis-acting elements had also been identified in different genes, including: the AuxRR-core and TGA-element (auxin-responsive element) (Ulmasov et al., 1997), the TCA-element (salicylic acid responsiveness; Zhang et al., 2017), the GARE-motif, and P-Box (gibberellin responsive element; Washida et al., 1999), the ERE (ethylene-responsive element; Fujimoto et al., 2000), suggesting *Shaker K^+^ channel* genes under regulated by different phytohormones and involved in corresponding plant physiological activity. Many cis-acting elements involved in plant stress response were also found on promotor region of Shaker K^+^ channel genes. Among them, MYB and MYC, two elements related to drought stress and abscisic acid-regulate (Abe et al., 1997), were predicted in all genes. And the MYC element was found in the largest number in Group II. The G-Box element was reported to be involved in chlorophyll synthesis (Abe et al., 1997), and was found to be the most abundant in *SiKAT3 (Seita.1G020100)*. The WUN-motif, a wound-responsive element (Hayashi et al., 2003), was only found in *SiAKT1 (Seita.5G244400)* and *SiGORK (Seita.7G111600)*. In conclusion, cis-acting element analysis showed that most of the Shaker K^+^ channel genes responded to different environmental stresses and different groups of genes might be under different regulate mechanisms.

**Figure 4.**
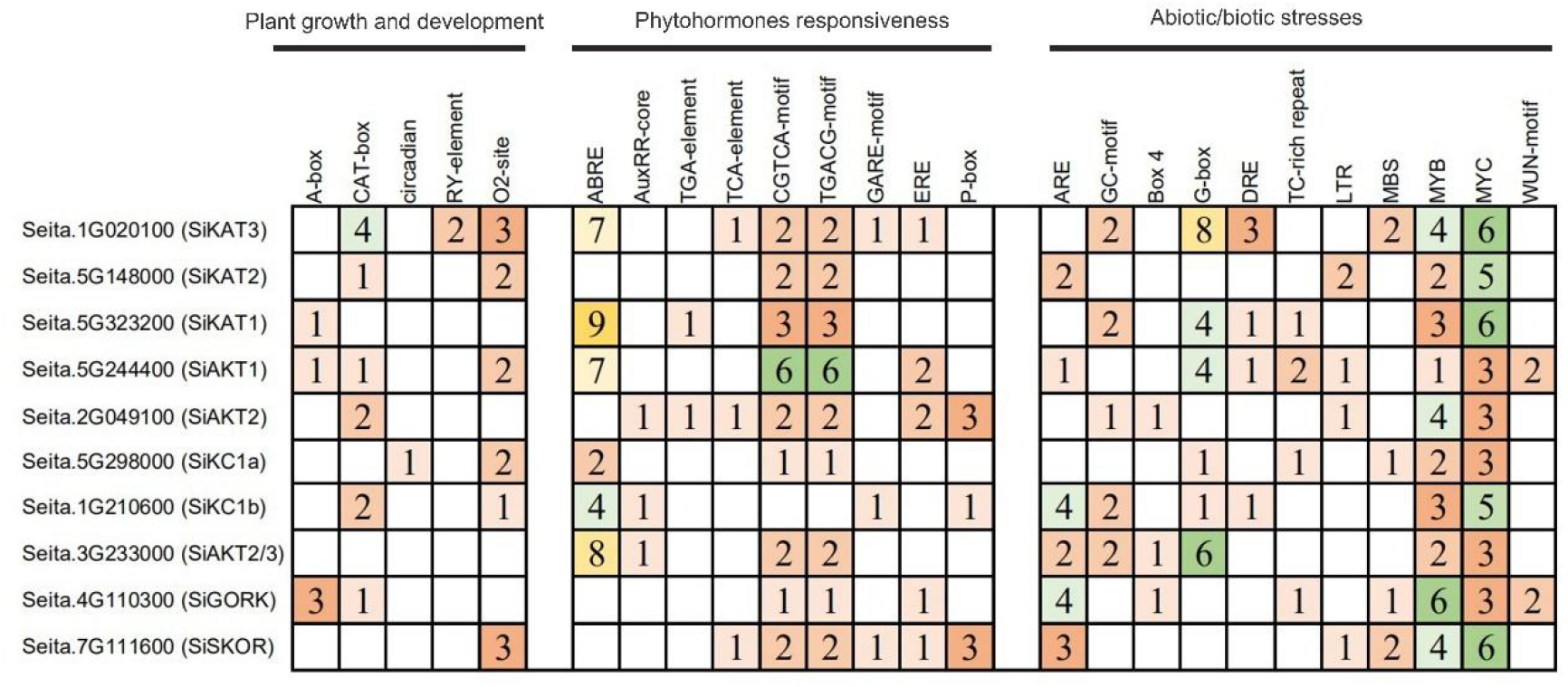
Cis-acting element analysis of the promoter regions of Shaker K^+^ channel genes from millet. The number of each cis-acting element in the promoter regions (2000 bp upstream of the translation start site) of *Shaker K^+^ channel* genes was shown in the figure. Based on the functional annotations, the cis-acting elements were classified into three major classes: plant growth and development, phytohormone responsiveness, and abiotic/biotic stresses related cis-acting elements.

### In silico transcript levels analysis of the Shaker K^+^ channel genes

To analyze the expression patterns of *Shaker K^+^ channel* genes in different tissues of millet, publicly available RNA-seq data was used (NCBI GEO RNA-seq DataSets GSE89855). This dataset including information from various tissues, including the inflorescence stage of root, the inflorescence and vegetative stages of shoot, the vegetative stage of four locations (tip, midsection, lower section, and base) of leaf, the inflorescence and vegetative stages of meristem, and the pre-anthesis and post-anthesis stages of panicle. The *SiAKT1 (Seita.5G244400)* was expressed highly in various tissues, excepted for meristem at the inflorescence stage and panicle at the pre-anthesis stage. The *SiAKT2/3 (Seita.3G233000)*was also expressed high in all tissues. The *SiKAT3 (Seita.1G020100)* could be found in all tissues above ground and the highest level was detected in leaf tip. Interestingly, according to this RNA-seq result, the *SiKAT1 (Seita.5G323200)* was not expressed in millet leaf and the two genes in group V *(SiGORK (Seita.7G111600)* and *SiSKOR (Seita.4G110300))* was only highly expressed in roots. The *SiAKT2 (Seita.2G0491000), SiKC1a (Seita.5G148000)*, and *SiKC1b (Seita.1G210600)* were almost not detected in all tissues.

### Prediction of Posttranslational modification sites on Shaker K^+^ channel genes from millet

Protein phosphorylation and asparagine (N)-linked glycosylation are two important post-translational modifications that affect the stability, subcellular location, and protein-protein interaction in plants (Xu et al., 2019). They are crucial for plant response to environment stresses (Jiao et al., 2020). As shown in Figure 6, these two post-translational modifications in Shaker K^+^ channel proteins from millet were predicted. SiKAT3 (Seita.1G020100) and SiAKT1 contain many glycosylation sites (five sites), SiKC1a, SiKC1b, and SiSKOR have three sites, SiAKT2 and SiAKT2/3 were predicted to have two N-glycosylation sites, and the remaining proteins only contained one site. Regarding the predicted phosphorylation sites, the Shaker K^+^channel proteins from millet contain a range from 40 to 79 (Figure 6 B). There were 79 sites in AKTs (SiAKT1 and SiAKT2), while in KC1s, 40 and 45 were predicted in SiKC1a and SiKC1b, respectively.

### Prediction of ABA binding site on SiGORK and SiSKOR

The plant hormone abscisic acid (ABA) plays an important role in plant response to abiotic stress, especially to drought stress (Fahad et al., 2015; Danquah et al., 2014). Previous report has shown that the GORK of Arabidopsis harbored potential ABA interaction sites (GORK^N558, K559, Y562, R565^) and the mutation of K559 and Y562 to Ala significantly reduced the outward K^+^ current of GORK, while the other two mutations of N558A/R565A resulted in the complete loss of GORK’s channel function (Ooi et al., 2017). As outward rectifying K^+^ channels, the relationship between SKOR and GORK is relatively close in evolution. Therefore, we compared the GORK and SKOR protein sequences of millet, Arabidopsis, tobacco, and other gramineous plants (rice, sorghum, wheat). As shown in Figure 6C, these four amino acid sites were existed in GORK protein of most species. Compared to the sequence of AtGORK, only N558 changes to Ser in SiGORK (Seita.7G111600), while the N558K559 changes to Lys-Asn in SiSKOR. Consider both Asn and Ser are polar amino acids with uncharged R groups, it is still possible that SiGORK contains potential ABA binding site like that of AtGORK.

### Expression analysis of Shaker K^+^ channel gene family under different abiotic stresses

To further study the role of Shaker K^+^ channel genes in plant stress resistance, fourteen days old seedlings of “Jingu21” was subjected to different abiotic treatments (cold, heat, NaCl, PEG, ABA). The seedlings did not show strong phenotype after 24 h treatment (Supplement Figure 1). Then the leaf samples were harvested at 0 h, 12 h, and 24 h after treatment. The expression patterns of these channel genes were detected by qPCR. The transcription level of four genes *(SiKAT1 (Seita.5G323200), SiAKT2 (Seita.2G049100), SiKC1a (Seita.5G148000), SiKC1b (Seita.1G210600))* were not detected because they were not expressed in the leaves, which was consistent with the result in Figure 5.

**Figure 5.**
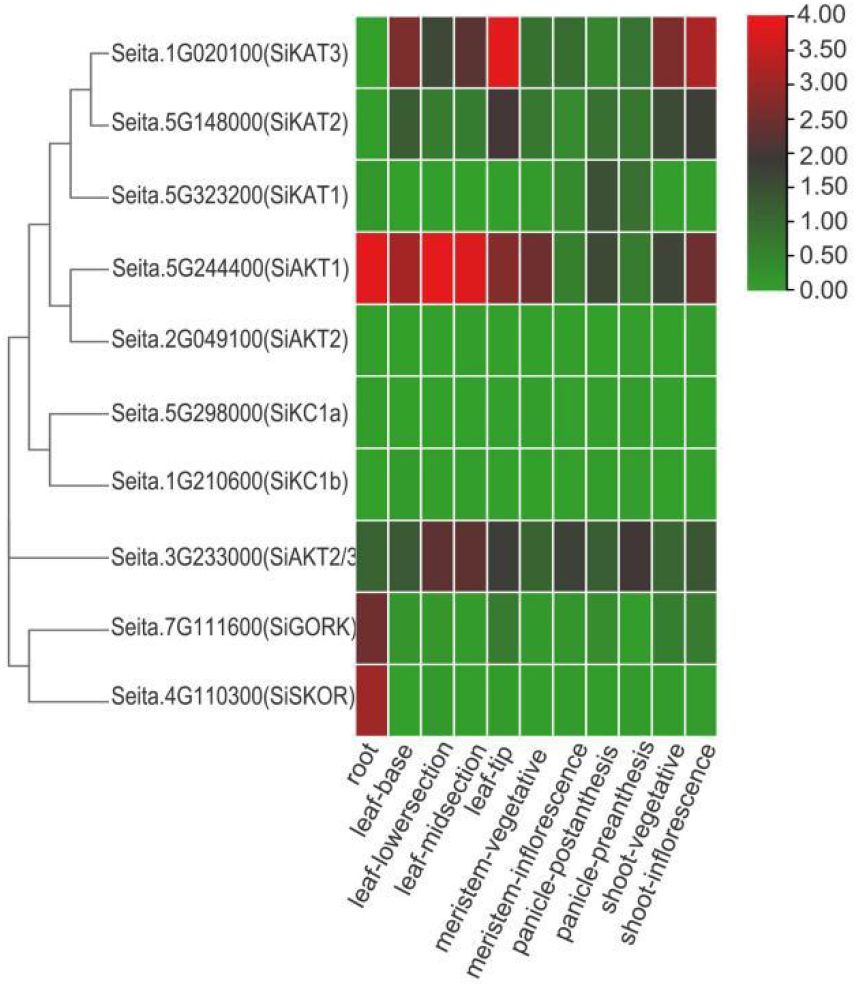
Expression profile of Shaker K^+^ channel genes in different tissues and developmental stages of millet. Gene expression data is downloaded from the NCBI GEO DataSets (GSE89855). Including roots (at inflorescence stage), shoot (at inflorescence and vegetative stages), leaf (four locations: tip, midsection, lower section, and base at the vegetative stage), meristem (at the inflorescence and vegetative stages), and panicle (at the pre-anthesis and postanthesis stages). The color scale represents expression levels (FPKM value) from high (red) to low (green colour).

**Figure 6.**
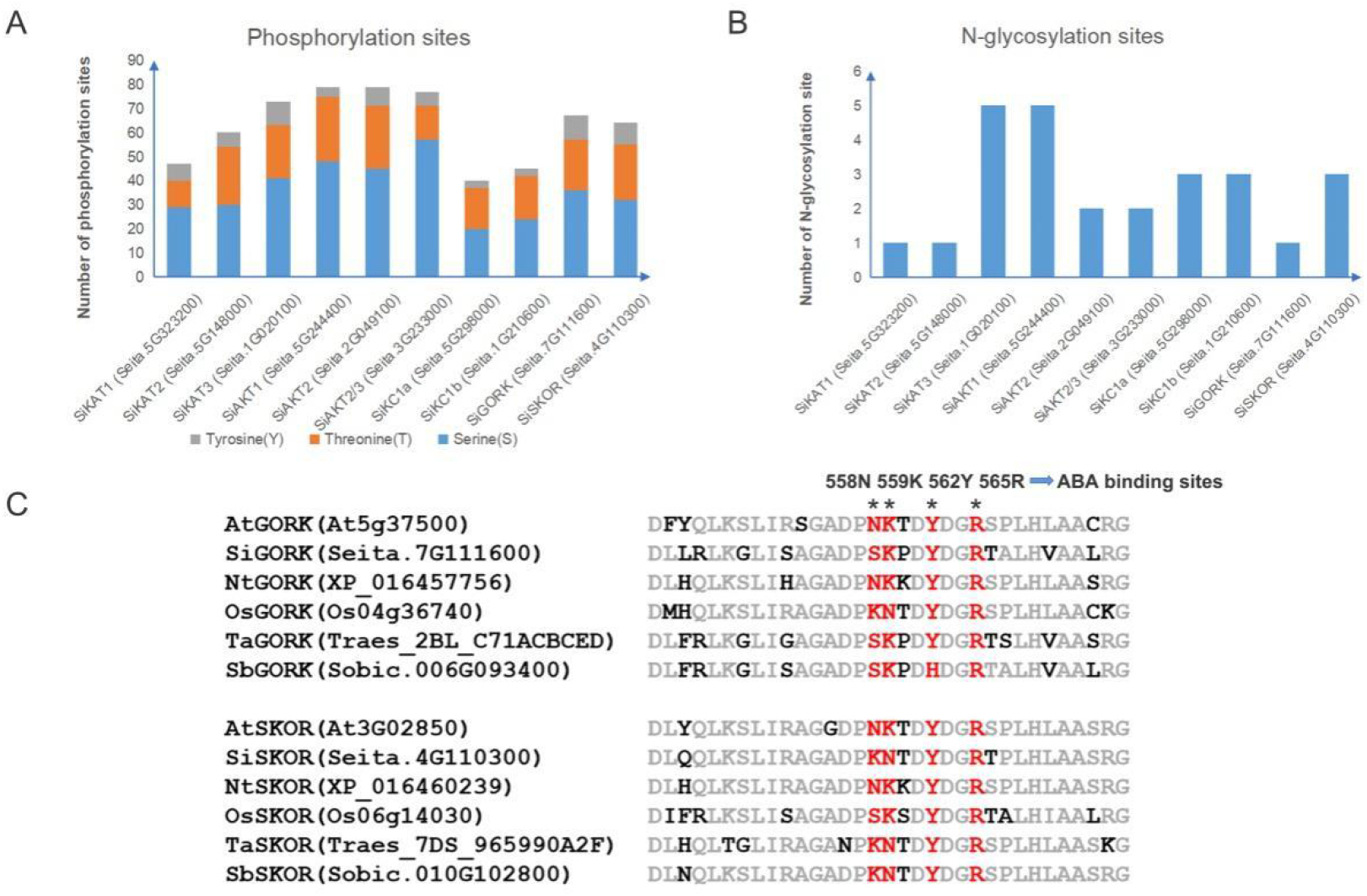
Prediction of posttranslational modification sites and ABA binding sites in the amino acid sequences of Shaker K^+^ channel proteins from millet. A: N-glycosylation site; B: phosphorylation site; C: ABA binding sites in GORK and SKOR

As shown in Figure 8, under cold stress (4°C), the transcript levels of *SiAKT1 (Seita.5G244400), SiAKT2/3 (Seita.3G233000)*, *SiKAT2 (Seita.5G148000), SiSKOR (Seita.4G110300)*, and *SiGORK (Seita.7G111600)* increased after 24 h treatment. For *SiSKOR (Seita.4G110300)*, its level was more significant up-regulated after 12 h cold treatment than that after 24 h cold treatment. Under hot stress (40°C day/32°C night), *SiAKT1 (Seita.5G244400)* was up-regulated after 24 h treatment. *SiSKOR (Seita.4G110300)* and *SiGORK (Seita.7G111600)* were increased strongly after 12 h treatment. The transcription of *SiAKT2/3 (Seita.3G233000)* was not responded to hot stress, while both cold and hot stress did not affect the level of *SiKAT3 (Seita.1G020100)*. These results suggest that most of the Shaker K^+^ channels from millet response differently to temperature stresses.

**Figure 8.**
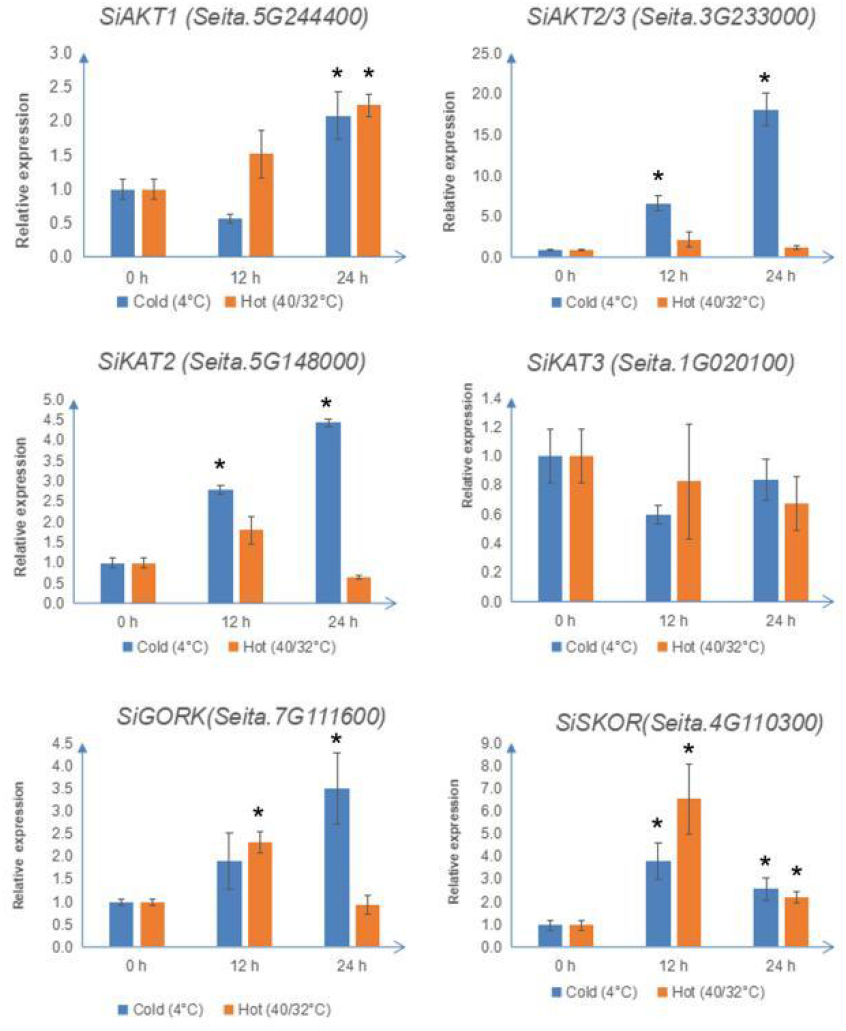
The relative transcript levels of Shaker K^+^ channel genes from millet under cold (4°C) or hot treatment (40°C day/32°C night). The qRT-PCR analyses were used to access transcript levels of Shaker K^+^ channel genes from millet with or without 12 h and 24 h cold (blue;4°C) or hot treatment (orange; 40 °C day/32°C night). Each bar represents the mean±SE normalized to *SiAct2 (Seita.8G043100)*. All samples were run in three biological and three technical replicates. Asterisk indicates that the gene expression after stress has a significant difference compared with the control at 0 h (*p < 0.05).

Further qRT-PCR analysis of millet Shaker K^+^ channel genes also showed their transcript levels varied under salt stress (150 mmol/l, 200 mmol/l) or PEG (10%, 15%) treatments. As shown in figure 9, transcription of *SiAKT1 (Seita.5G244400)* decreased under both treatments compared with the control. *SiAKT2 (Seita.3G233000)* was up-regulated after 24 h 10% and 15% PEG treatment. For *SiKAT2 (Seita.5G148000)*, 200 mmol/l salt stress slightly induced its transcription at 12 h; 10% PEG treatment decreased its expression at 12 h, but significantly up-regulated it after 24 h. At 12 h, *SiKAT3 (Seita.1G020100)* was strongly induced only by 10% PEG treatment. At 24 h, this gene was induced by both 10% and 15% PEG treatment. For outward rectifying channels, *SiGORK (Seita.7G111600)* decreased under PEG treatment in both two time points, while *SiSKOR (Seita.4G110300)* was up-regulated by salt stress and 10 % PEG treatment after 24 h.

**Figure 9.**
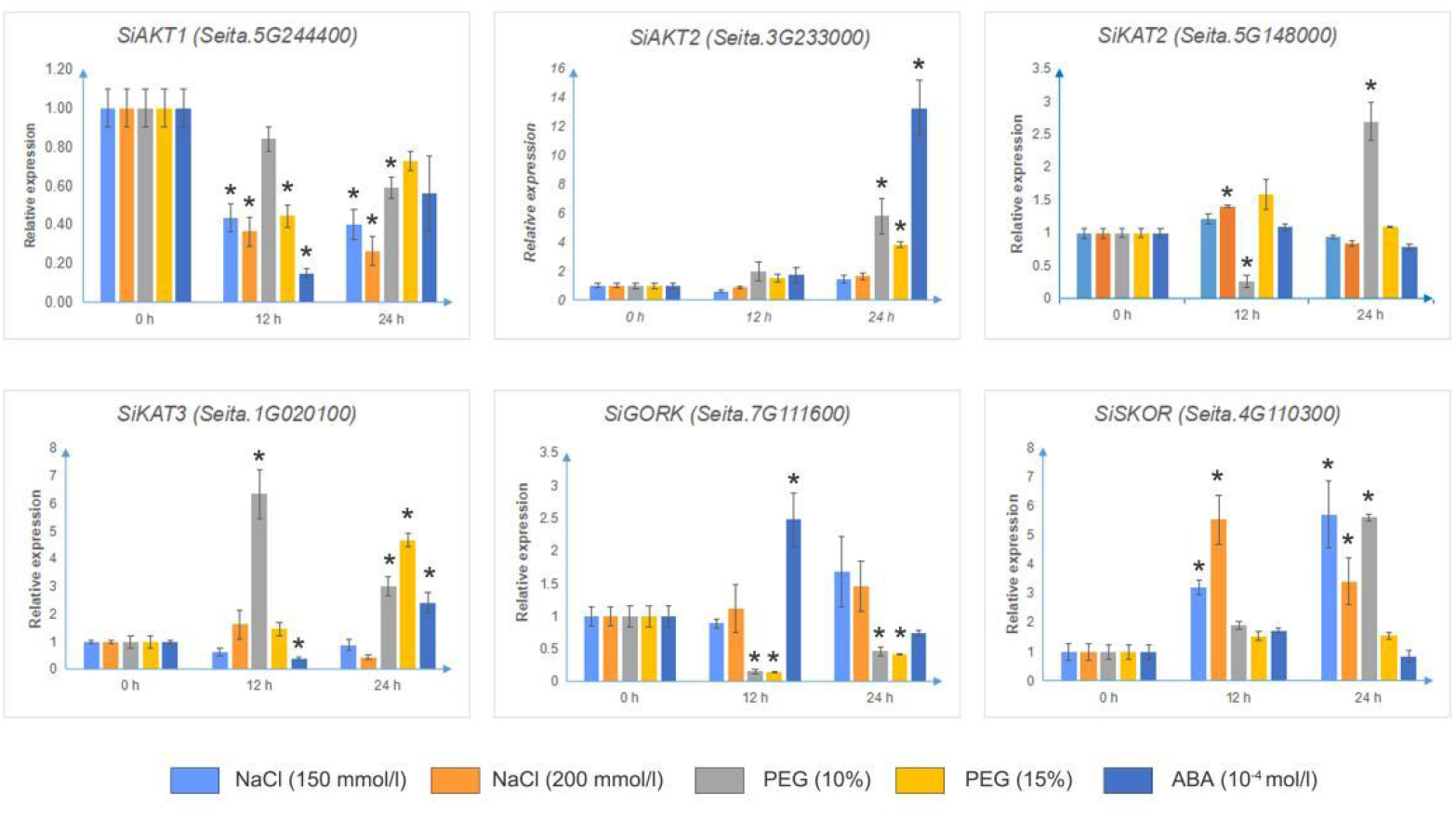
The relative transcript levels of Shaker K^+^ channel genes from millet under different stress. The qRT-PCR analyses were used to access transcript levels of Shaker K^+^ channel genes from millet with or without 12 h and 24 h NaCl (150 or 200 mmol/l), PEG (10% or 15%), or ABA (10^-4^ mol/l). Each bar represents the mean ±SE normalized to *SiAct2 (Seita.8G043100)*. All samples were run in three biological and three technical replicates. Asterisk indicates that the gene expression after stress has a significant difference compared with the control at 0 h (*p < 0.05).

ABA (10 mmol/l) treatment decreased the transcription of *SiAKT1 (Seita.5G244400)* only at 12 h. The expression of *SiAKT2 (Seita.3G233000)* and *SiKAT3 (Seita.1G020100)* increased significantly after 24 hours of treatment. Especially for *SiKAT3*, its transcript was up-regulated 10 times compared to the control. The expression of other Shaker K^+^ channel genes showed no change under ABA treatment. Most detectable channel genes were induced by either codl or hot stress.

## Discussion

Potassium (K^+^) is essential for plant growth and involved in stress resistance in higher plants (Wang et al., 2013). Shaker K^+^ channels are highly conserved voltage-dependent ion channels in plants, which play important role in the transport and distribution of K^+^ in plant cells. Investigations focused on this family of genes have been performed in many species (Dreyer and Uozumi, 2011; Amrutha et al., 2007; Bauer et al., 2000; Jin et al., 2021). However, these is little report focused on the Shaker K^+^ channels in foxtail millet (*Setaria italica*), a high drought resistance cereal crops wildly cultivated in Asia.

In the current study, we identified ten members belong to the Shaker K^+^ channel family from millet based on the gene information from rice and Arabidopsis. Previous research suggested that Shaker K^+^ channels are highly conserved in plants (Dreyer and Blatt, 2009). These genes from millet were classified into five different groups as reported in other species (Table 1 and Figure 1). Further analysis showed that channel genes of each group shared similar motifs and gene structures (Figure 3). Based on the phylogenetic results, the channel genes from millet were more similar to their orthologous genes in rice than in Arabidopsis, suggesting the diversification of Shaker K^+^ channels happened before the separation of monocots and dicots in evolution, supporting previous report (Pilot et al., 2003b). The five-group classification of channels is based on their functional diversity (Dreyer and Blatt, 2009; Pilot et al., 2003b), so we can predict the channel activity of each group in millet depends on their classification. In millet, there were two members in group IV in millet (SiKC1a and SiKC1b), while there were only one in Arabidopsis and three in rice. KC1s in group IV is silent when expressed alone, but function as general regulator of channel activity when form hetero-tetramer with other channel proteins (Jeanguenin et al., 2011; Duby et al., 2008). It is possible that the function provided by channels from Group IV is important for each species adapted to their specific environment condition.

To identify the role of Shaker K^+^ channels in abiotic stress response, we first analyzed the Cis-acting element in promotor region of each gene. As shown in figure 4, various types of cis-acting elements existed in Shaker K^+^ channel genes, even in the same group, suggesting these genes express differently in response to different environment stresses in millet. Cis-acting element analysis of channel genes revealed the exist of various hormone response elements, including ABA, auxin, GA, MeJA, and ethylene. The MYB and MYC elements were identified in most of the genes, suggesting that Shaker K^+^ channel genes in millet were regulated by drought stress, which needs to be experimentally proved.

Expression profiling using previously released RNA-seq data (NCBI GEO RNA-seq DataSets GSE89855) was also performed. Different Shaker K^+^ channel genes showed differential expression levels in different tissues. SiAKT1 expressed in all tissues, especially highly expressed in root, supporting its role in K^+^ absorbing from soil as its orthologous genes in other species (Véry and Sentenac, 2003; Li et al., 2014). SiKAT3 was highly expressed in shoot and leaf, suggesting its role might like AtKAT1, which involved in the regulation of stomata movement (Wang and Wu, 2013). Interestingly, according to the RNA-seq data, the genes in group IV could not detect in all tissues. One explanation of this result is that the expression of some K^+^ channels are only induced under stress (Zhang et al., 2018; Jin et al., 2021), which need further research.

Glycosylation and phosphorylation modifications play vital roles in protein functions in plant. The potential posttranslational modifications of Shaker K^+^ channel proteins in millet were also predicted. Our results showed that AKTs like SiAKT1 and SiAKT2 had largest number of modification sites (Figure 6). Previous reports have shown that AtAKT1 was activated by phosphorylation, which depends on a calcineurin B-like 1 (CBL1) with a CBL-interacting protein kinase (CIPK 23) (Xu et al., 2006; Honsbein et al., 2009). Such phosphorylation activating might also happened in other Shaker K^+^ channels in millet.

As mentioned above, to investigate whether Shaker K^+^ channels in millet were regulated by abiotic stress or ABA treatment and whether SiKC1s will be induced under stress, we performed quantitative real-time PCR analysis in fourteen days old seedlings of “Jingu21” under different abiotic stress treatments (cold, heat, NaCl, PEG) and ABA treatment. “Jingu21” was one of the cultivars produced by radiation induced mutagenesis in breeding program (He et al., 2015). This cultivar shows good quality, high yield, and better drought stress-tolerant (Wang et al., 2021). Our results showed that the transcription level of four Shaker K^+^ channel genes (*SiKAT1*, *SiAKT2*, *SiKC1a*, and *SiKC1b*) were not detected, even under different abiotic stress and ABA treatment. This finding support RNA-seq data shown in Figure 5. However, as important regulator channel proteins, the missing of SiKC1s left a question mark in the present study. *AtKC1* was upregulated by salt stress and K^+^ deficiency (Pilot et al., 2003a). The transcription levels of SiKC1s under low K^+^ stress was worth to test in future research. AtKC1 and AtAKT1 form functional channel in root (Geiger et al., 2009; Duby et al., 2008). Whether SiAKT1 forms functional channel with other Shaker K^+^ channel proteins in millet might be one other explanation for the question of SiKC1s here. SiAKT2/3 also found in most tissues (Figure 5). It is worth to co-express SiAKT2/3 with SiAKT1 and detected the inward K^+^ current change by the electrophysiological experiment.

As shown in Figure 8 and Figure 9, other remaining gene displayed its own regulation pattern under different treatments, which was consistent with findings from previous reports in Arabidopsis (Pilot et al., 2003a) and sweetpotato (Jin et al., 2021). In Arabidopsis, AKT1 is involved in a major pathway for K^+^ uptake at the root epidermis (Geiger et al., 2009). SiAKT1 induced by temperature stress and decreased under salt and osmotic stress revealed the role of K^+^ absorption in plant adaptation to different environment conditions. SiSKOR might be involved in the K^+^ secretion into the xylem sap as its orthologous gene in Arabidopsis (Dreyer and Blatt, 2009; Johansson et al., 2006). Our qRT-PCR analysis revealed that this outward K^+^channel was upregulated by all stress and un-response to ABA in the present study. Previous reports suggested that SKOR was alter regulated by different abiotic stress and decreased by ABA treatment (Pilot et al., 2003a; Jin et al., 2021). The special pattern of *SiSKOR* compared to SKORs in other species suggest the redistribution of K^+^ among tissues might contribute to the stress tolerant of millet. Previous report has shown that ABA does not affect the transcription level of GORK, but ABA induced plasma membrane depolarization and activate GORK in guard cell, leading to stomata closure (Hosy et al., 2003; Pilot et al., 2003a; Thiel et al., 1992). Except *SiAKT2* and *SiKAT3*, we found ABA treatment had little or no effect on the transcription of other Shaker K^+^ channels in millet. In 2017, Ooi et al. reported that there is direct GORK–ABA interaction which enhance K^+^-efflux current through GORK (Ooi et al., 2017). The most important sits for ABA docking are N558, K559, Y562, and R565 at the C terminal of GORK. So, we performed a protein sequence alignment among GORK and SKOR from different species. We found such sites were existed on most species, especially on channel of millet (Figure 6C), suggesting these outward channels in millet might under regulation of ABA through directly binding.

In conclusion, we identified ten Shaker K^+^ channel genes in Foxtail millet and performed a basic analysis for their role in plant stress response through cis-acting element analysis, RNA-Seq data, and qRT-PCR analysis. This work will facilitate further research focused on the biological roles of Shaker K^+^ genes in millet.

## Declarations

Conflict of interests All authors have no relevant financial or non-financial interests to disclose.

## Authors’ contributions

Experiments were designed by Ben Zhang, Lizhen Zhang, Pu Yang, and Hui Wang. Experiments were performed by Hui Wang, Yue Guo, Xiaoxia Wang, and Mengtao Lv. Ben Zhang, Hui Wang, and Lizhen Zhang analyzed the data and wrote the manuscript. All authors read and approved the final manuscript.

## Data availability

The data and materials generated and/or analyzed during the current study are available from the corresponding author on reasonable request.

## Supplement information

**Supplement Figure 1.**
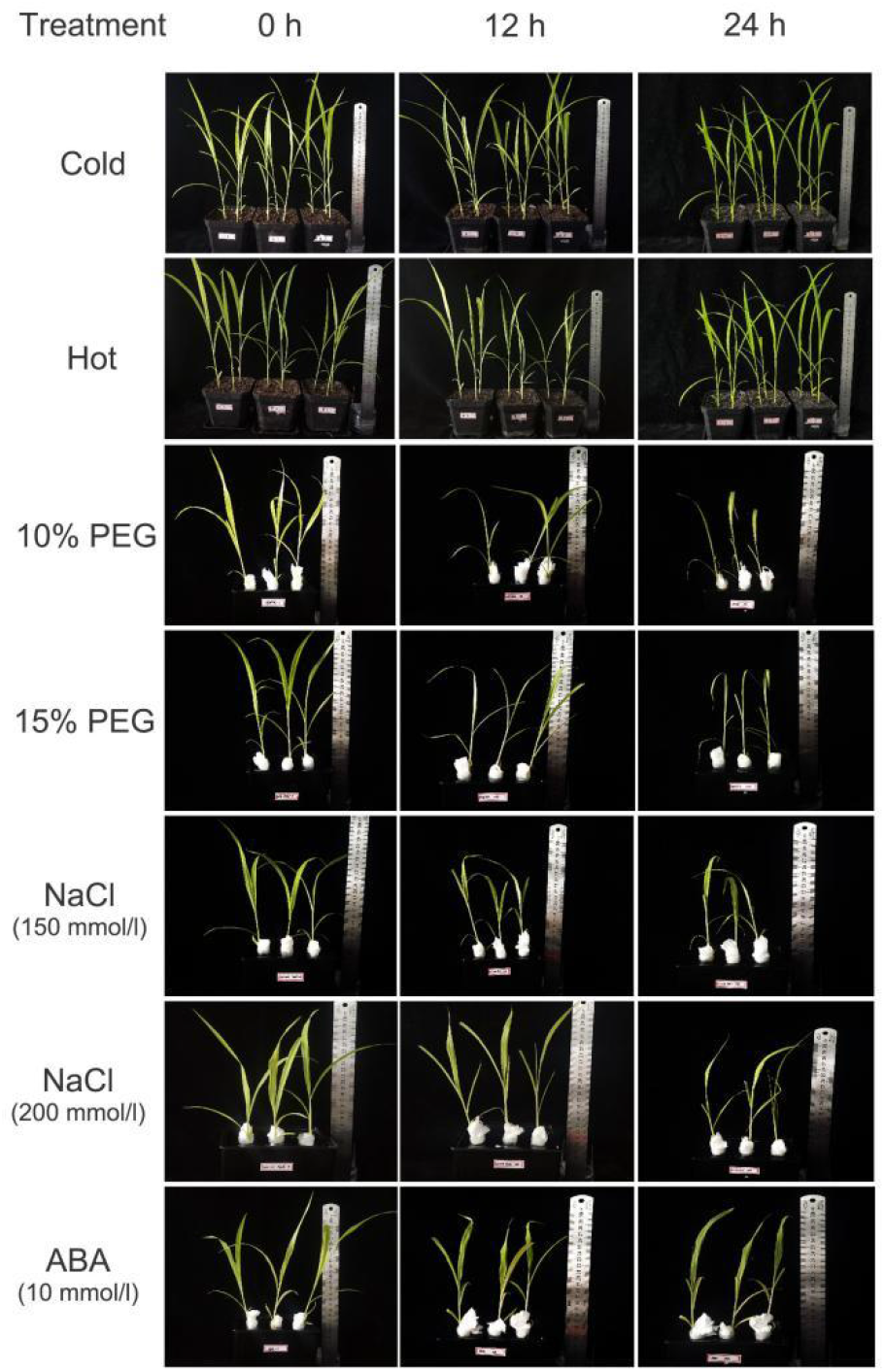

## References

Abe, H., Yamaguchi-Shinozaki, K., Urao, T., Iwasaki, T., Hosokawa, D., and Shinozaki, K. (1997). Role of Arabidopsis MYC and MYB homologs in drought- and abscisic acid-regulated gene expression. Plant Cell 9: 1859–1868.

Ahmad, I., Mian, A., and Maathuis, F.J.M. (2016). Overexpression of the rice AKT1 potassium channel affects potassium nutrition and rice drought tolerance. J. Exp. Bot. 67: 2689–2698.

Amrutha, R.N., Sekhar, P.N., Varshney, R.K., and Kishor, P.B.K. (2007). Genome-wide analysis and identification of genes related to potassium transporter families in rice (Oryza sativa L.). Plant Sci. 172: 708–721.

Anderson, S.L., Teakle, G.R., Martino-Catt, S.J., and Kay, S.A. (1994). Circadian clock- and phytochrome-regulated transcription is conferred by a 78 bp cis-acting domain of the Arabidopsis CAB2 promoter. Plant J. 6: 457–470.

Bailey, T.L., Boden, M., Buske, F.A., Frith, M., Grant, C.E., Clementi, L., Ren, J., Li, W.W., and Noble, W.S. (2009). MEME Suite: Tools for motif discovery and searching. Nucleic Acids Res. 37: 202–208.

Bassham, D.C. and Blatt, M.R. (2008). SNAREs: cogs and coordinators in signaling and development. Plant Physiol. 147: 1504–1515.

Bauer, C.S., Hoth, S., Haga, K., Philippar, K., Aoki, N., and Hedrich, R. (2000). Differential expression and regulation of K+ channels in the maize coleoptile: Molecular and biophysical analysis of cells isolated from cortex and vasculature. Plant J. 24: 139–145.

Becker, D., Hoth, S., Ache, P., Wenkel, S., Roelfsema, M.R.G., Meyerhoff, O., Hartung, W., and Hedrich, R. (2003). Regulation of the ABA-sensitive Arabidopsis potassium channel gene GORK in response to water stress. FEBS Lett. 554: 119–126.

Bennetzen, J.L. et al. (2012). Reference genome sequence of the model plant Setaria. Nat. Biotechnol. 30: 555–561.

Cao, Y., Ward, J.M., Kelly, W.B., Ichida, A.M., Gaber, R.F., Anderson, J.A., Uozumi, N., Schroeder, J.I., and Crawford, N.M. (1995). Multiple genes, tissue specificity, and expression-dependent modulation contribute to the functional diversity of potassium channels in Arabidopsis thaliana. Plant Physiol. 109: 1093–1106.

Chen, C., Chen, H., Zhang, Y., Thomas, H.R., Frank, M.H., He, Y., and Xia, R. (2020). TBtools: An Integrative Toolkit Developed for Interactive Analyses of Big Biological Data. Mol. Plant 13: 1194–1202.

Chen, X., Ding, Y., Yang, Y., Song, C., Wang, B., Yang, S., Guo, Y., and Gong, Z. (2021). Protein kinases in plant responses to drought, salt, and cold stress. J. Integr. Plant Biol. 63: 53–78.

Chen, Z., Zhou, M., Newman, I.A., Mendham, N.J., Zhang, G., and Shabala, S. (2007). Potassium and sodium relations in salinised barley tissues as a basis of differential salt tolerance. Funct. Plant Biol. 34: 150–162.

Danquah, A., de Zelicourt, A., Colcombet, J., and Hirt, H. (2014). The role of ABA and MAPK signaling pathways in plant abiotic stress responses. Biotechnol. Adv. 32: 40–52.

Dreyer, I. and Blatt, M.R. (2009). What makes a gate? The ins and outs of Kv-like K+ channels in plants. Trends Plant Sci. 14: 383–390.

Dreyer, I. and Uozumi, N. (2011). Potassium channels in plant cells. FEBS J. 278: 4293–4303.

Duby, G., Hosy, E., Fizames, C., Alcon, C., Costa, A., Sentenac, H., and Thibaud, J.B. (2008). AtKC1, a conditionally targeted Shaker-type subunit, regulates the activity of plant K+ channels. Plant J. 53: 115–123.

Fahad, S. et al. (2015). Phytohormones and plant responses to salinity stress: a review. Plant Growth Regul. 75: 391–404.

Feng, X., Liu, W., Cao, F., Wang, Y., Zhang, G., Chen, Z.-H., and Wu, F. (2020). Overexpression of HvAKT1 improves drought tolerance in barley by regulating root ion homeostasis and ROS and NO signaling. J. Exp. Bot. 71: 6587–6600.

Fujimoto, S.Y., Ohta, M., Usui, A., Shinshi, H., and Ohme-Takagi, M. (2000). Arabidopsis ethylene-responsive element binding factors act as transcriptional activators or repressors of GCC box-mediated gene expression. Plant Cell 12: 393–404.

Gaber, R.F., Styles, C.A., and Fink, G.R. (1988). TRK1 encodes a plasma membrane protein required for high-affinity potassium transport in Saccharomyces cerevisiae. Mol. Cell. Biol. 8: 2848–2859.

Gasteiger, E., Gattiker, A., Hoogland, C., Ivanyi, I., Appel, R.D., and Bairoch, A. (2003). ExPASy: The proteomics server for in-depth protein knowledge and analysis. Nucleic Acids Res. 31: 3784–3788.

Geiger, D., Becker, D., Vosloh, D., Gambale, F., Palme, K., Rehers, M., Anschuetz, U., Dreyer, I., Kudla, J., and Hedrich, R. (2009). Heteromeric AtKC1·AKT1 channels in Arabidopsis roots facilitate growth under K+-limiting conditions. J. Biol. Chem. 284: 21288–21295.

Gong, Z., Xiong, L., Shi, H., Yang, S., Herrera-estrella, L.R., Xu, G., Chao, D., Li, J., and Wang, P. (2020). Plant abiotic stress response and nutrient use efficiency. Sci. CHINA Life Sci. 336: 1–40.

Goodstein, D.M., Shu, S., Howson, R., Neupane, R., Hayes, R.D., Fazo, J., Mitros, T., Dirks, W., Hellsten, U., Putnam, N., and Rokhsar, D.S. (2012). Phytozome: A comparative platform for green plant genomics. Nucleic Acids Res. 40: 1178–1186.

Grefen, C., Obrdlik, P., and Harter, K. (2009). The determination of protein-protein interactions by the mating-based split-ubiquitin system (mbSUS). In Methods in Molecular Biology, pp. 217–233.

Guerriero, G., Martin, N., Golovko, A., Sundström, J.F., Rask, L., and Ezcurra, I. (2009). The RY/Sph element mediates transcriptional repression of maturation genes from late maturation to early seedling growth. New Phytol. 184: 552–565.

Hayashi, T., Kobayashi, D., Kariu, T., Tahara, M., Hada, K., Kouzuma, Y., and Kimura, M. (2003). Genomic cloning of ribonucleases in Nicotiana glutinosa leaves, as induced in response to wounding or to TMV-infection, and characterization of their promoters. Biosci. Biotechnol. Biochem. 67: 2574–2583.

He, L., Zhang, B., Wang, X., Li, H., and Han, Y. (2015). Foxtail millet: Nutritional and eating quality, and prospects for genetic improvement. Front. Agric. Sci. Eng. 2: 124–133.

Hobo, T., Asada, M., Kowyama, Y., and Hattori, T. (1999). ACGT-containing abscisic acid response element (ABRE) and coupling element 3 (CE3) are functionally equivalent. Plant J. 19: 679–689.

Honsbein, A., Sokolovski, S., Grefen, C., Campanoni, P., Pratelli, R., Paneque, M., Chen, Z., Johansson, I., and Blatt, M.R. (2009). A tripartite SNARE-K+ channel complex mediates in channel-dependent K+ nutrition in Arabidopsis. Plant Cell 21: 2859–2877.

Horaruang, W. and Zhang, B. (2017). Mating Based Split-ubiquitin Assay for Detection of Protein Interactions. Bio-Protocol 7: 1–14.

Horton, P., Park, K.J., Obayashi, T., Fujita, N., Harada, H., Adams-Collier, C.J., and Nakai, K. (2007). WoLF PSORT: Protein localization predictor. Nucleic Acids Res. 35: 585–587.

Hosy, E. et al. (2003). The Arabidopsis outward K+ channel GORK is involved in regulation of stomatal movements and plant transpiration. Proc. Natl. Acad. Sci. U. S. A. 100: 5549–5554.

Hu, B., Jin, J., Guo, A.Y., Zhang, H., Luo, J., and Gao, G. (2015). GSDS 2.0: An upgraded gene feature visualization server. Bioinformatics 31: 1296–1297.

Hwang, H., Yoon, J., Kim, H.Y., Min, M.K., Kim, J.A., Choi, E.H., Lan, W., Bae, Y.M., Luan, S., Cho, H., and Kim, B.G. (2013). Unique Features of Two Potassium Channels, OsKAT2 and OsKAT3, Expressed in Rice Guard Cells. PLoS One 8: 1–14.

Jeanguenin, L., Alcon, C., Duby, G., Boeglin, M., Chérel, I., Gaillard, I., Zimmermann, S., Sentenac, H., and Véry, A.A. (2011). AtKC1 is a general modulator of Arabidopsis inward Shaker channel activity. Plant J. 67: 570–582.

Jeanguenin, L., Lebaudy, A., Xicluna, J., Alcon, C., Hosy, E., Duby, G., Michard, E., Lacombe, B., Dreyer, I., and Thibaud, J.B. (2008). Heteromerization of Arabidopsis Kv channel α-subunits: Data and prospects. Plant Signal. Behav. 3: 622–625.

Jiao, Q.S., Niu, G.T., Wang, F.F., Dong, J.Y., Chen, T.S., Zhou, C.F., and Hong, Z. (2020). N-glycosylation regulates photosynthetic efficiency of arabidopsis thaliana. Photosynthetica 58: 72–79.

Jin, R., Zhang, A., Sun, J., Chen, X., Liu, M., Zhao, P., Jiang, W., and Tang, Z. (2021). Identification of Shaker K+ channel family members in sweetpotato and functional exploration of IbAKT1. Gene 768: 145311.

Johansson, I., Wulfetange, K., Porée, F., Michard, E., Gajdanowicz, P., Lacombe, B., Sentenac, H., Thibaud, J.B., Mueller-Roeber, B., Blatt, M.R., and Dreyer, I. (2006). External K+ modulates the activity of the Arabidopsis potassium channel SKOR via an unusual mechanism. Plant J. 46: 269–281.

Kumar, S., Stecher, G., and Tamura, K. (2016). MEGA7: Molecular Evolutionary Genetics Analysis Version 7.0 for Bigger Datasets. Mol. Biol. Evol. 33: 1870–1874.

Kwon, C., Lee, J., and Yun, H.S. (2020). SNAREs in Plant Biotic and Abiotic Stress Responses. Mol. Cells 25: 501–508.

Lebaudy, A., Vavasseur, A., Hosy, E., Dreyer, I., Leonhardt, N., Thibaud, J.B., Véry, A.A., Simonneau, T., and Sentenac, H. (2008). Plant adaptation to fluctuating environment and biomass production are strongly dependent on guard cell potassium channels. Proc. Natl. Acad. Sci. U. S. A. 105: 5271–5276.

Lebaudy, A., Véry, A.A., and Sentenac, H. (2007). K+ channel activity in plants: Genes, regulations and functions. FEBS Lett. 581: 2357–2366.

Lelievre, J.M., Oliveira, L.O., and Nielsen, N.C. (1992). 5’-CATGCAT-3’ elements modulate the expression of glycinin genes. Plant Physiol. 98: 387–391.

Lescot, M., Déhais, P., Thijs, G., Marchal, K., Moreau, Y., Van De Peer, Y., Rouzé, P., and Rombauts, S. (2002). PlantCARE, a database of plant cis-acting regulatory elements and a portal to tools for in silico analysis of promoter sequences. Nucleic Acids Res. 30: 325–327.

Li, J., Yu, L., Qi, G.N., Li, J., Xu, Z.J., Wu, W.H., and Yi, W. (2014). The Os-AKT1 channel is critical for K+ uptake in rice roots and is modulated by the rice CBL1-CIPK23 complex. Plant Cell 26: 3387–3402.

Ma, W., Yang, G., Xiao, Y., Zhao, X., and Wang, J. (2020). ABA-dependent K+ flux is one of the important features of the drought response that distinguishes Catalpa from two different habitats. Plant Signal. Behav. 15.

Ooi, A., Lemtiri-Chlieh, F., Wong, A., and Gehring, C. (2017). Direct Modulation of the Guard Cell Outward-Rectifying Potassium Channel (GORK) by Abscisic Acid. Mol. Plant 10: 1469–1472.

Peng, R. and Zhang, B. (2020). Foxtail Millet: A New Model for C4 Plants. Trends Plant Sci.: 3–5.

Pilot, G., Gaymard, F., Mouline, K., Chérel, I., and Sentenac, H. (2003a). Regulated expression of Arabidopsis Shaker K+ channel genes involved in K+ uptake and distribution in the plant. Plant Mol. Biol. 51: 773–787.

Pilot, G., Pratelli, R., Gaymard, F., Meyer, Y., and Sentenac, H. (2003b). Five-group distribution of the Shaker-like K+ channel family in higher plants. J. Mol. Evol. 56: 418–434.

Raddatz, N., Morales de los Ríos, L., Lindahl, M., Quintero, F.J., and Pardo, J.M. (2020). Coordinated Transport of Nitrate, Potassium, and Sodium. Front. Plant Sci. 11: 1–18.

Salinas-cornejo, J., Madrid-espinoza, J., and Ruiz-lara, S. (2019). Identification and transcriptional analysis of SNARE vesicle fusion regulators in tomato (Solanum lycopersicum) during plant development and a comparative analysis of the response to salt stress with wild relatives. J. Plant Physiol. 242: 153018.

Sutter, J.-U., Campanoni, P., Tyrrell, M., and Blatt, M.R. (2006). Selective mobility and sensitivity to SNAREs is exhibited by the Arabidopsis KAT1 K+ channel at the plasma membrane. Plant Cell 18: 935–954.

Sutter, J.U., Sieben, C., Hartel, A., Eisenach, C., Thiel, G., and Blatt, M.R. (2007). Abscisic Acid Triggers the Endocytosis of the Arabidopsis KAT1 K+ Channel and Its Recycling to the Plasma Membrane. Curr. Biol. 17: 1396–1402.

Thiel, G., MacRobbie, E.A.C., and Blatt, M.R. (1992). Membrane transport in stomatal guard cells: The importance of voltage control. J. Membr. Biol. 126: 1–18.

Thompson, J.D., Gibson, T.J., and Higgins, D.G. (2003). Multiple Sequence Alignment Using ClustalW and ClustalX. Curr. Protoc. Bioinforma. 00: 1–22.

Ulmasov, T., Murfett, J., Hagen, G., and Guilfoyle, T.J. (1997). Creation of a Highly Active Synthetic AuxRE. Society 9: 1963–1971.

Véry, A.-A. and Sentenac, H. (2003). M Olecular M Echanisms and R Egulation of K + T Ransport in H Igher P Lants. Annu. Rev. Plant Biol. 54: 575–603.

Wang, H., Hao, D., Wang, X., Zhang, H., Yang, P., Zhang, L., and Zhang, B. (2021). Genome - wide identification and expression analysis of the SNARE genes in Foxtail millet (Setaria italica) reveals its roles in drought stress. Plant Growth Regul.

Wang, M., Zheng, Q., Shen, Q., and Guo, S. (2013). The critical role of potassium in plant stress response. Int. J. Mol. Sci. 14: 7370–7390.

Wang, Y., Salasini, B.C., Khan, M., Devi, B., Bush, M., Subramaniam, R., and Hepworth, S.R. (2019). Clade i tgacg-motif binding basic leucine zipper transcription factors mediate blade-on-petiole-dependent regulation of development. Plant Physiol. 180: 937–951.

Wang, Y. and Wu, W.H. (2013). Potassium transport and signaling in higher plants. Annu. Rev. Plant Biol. 64: 451–476.

Ward, J.M., Mäser, P., and Schroeder, J.I. (2009). Plant ion channels. Annu Rev Physiol 71: 59–82.

Washida, H., Wu, C.Y., Suzuki, A., Yamanouchi, U., Akihama, T., Harada, K., and Takaiwa, F. (1999). Identification of cis-regulatory elements required for endosperm expression of the rice storage protein glutelin gene GluB-1. Plant Mol. Biol. 40: 1–12.

Wu, G.Q., Li, Z.Q., Cao, H., and Wang, J.L. (2019). Genome-wide identification and expression analysis of the WRKY genes in sugar beet (Beta vulgaris L.) under alkaline stress. PeerJ 2019.

Xu, J., Li, H.D., Chen, L.Q., Wang, Y., Liu, L.L., He, L., and Wu, W.H. (2006). A Protein Kinase, Interacting with Two Calcineurin B-like Proteins, Regulates K+ Transporter AKT1 in Arabidopsis. Cell 125: 1347–1360.

Xu, S., Xiao, J., Yin, F., Guo, X., Xing, L., Xu, Y., and Chong, K. (2019). The protein modifications of O-GlcNAcylation and phosphorylation mediate vernalization response for flowering in winter wheat. Plant Physiol. 180: 1436–1449.

Yu, C.-S., Lin, C.-J., and Hwang, J.-K. (2004). Predicting subcellular localization of proteins for Gram-negative bacteria by support vector machines based on n -peptide compositions . Protein Sci. 13: 1402–1406.

Zhang, B., Karnik, R., Wang, Y., Wallmeroth, N., Blatt, M.R., and Grefen, C. (2015). The Arabidopsis R-SNARE VAMP721 Interacts with KAT1 and KC1 K + Channels to Moderate K + Current at the Plasma Membrane. Plant Cell 27: 1697–1717.

Zhang, B., Wang, H., and Zhang, Y. (2020). SNARE proteins and their role in plant ion channel regulation. Plant Growth Regul.

Zhang, G. et al. (2012). Genome sequence of foxtail millet (Setaria italica) provides insights into grass evolution and biofuel potential. Nat. Biotechnol. 30: 549–554.

Zhang, H., Xiao, W., Yu, W., Yao, L., Li, L., Wei, J., and Li, R. (2018). Foxtail millet SiHAK1 excites extreme high-affinity K+ uptake to maintain K+ homeostasis under low K+ or salt stress. Plant Cell Rep. 37: 1533–1546.

Zhang, H., Yin, W., and Xia, X. (2010). Shaker-like potassium channels in Populus, regulated by the CBL-CIPK signal transduction pathway, increase tolerance to low-K+ stress. Plant Cell Rep. 29: 1007–1012.

Zhang, R.X., Qin, L.J., and Zhao, D.G. (2017). Overexpression of the OsIMP gene increases the accumulation of inositol and confers enhanced cold tolerance in tobacco through modulation of the antioxidant enzymes’ activities. Genes (Basel). 8.

